# Structural basis for HKU5‑CoV main protease inhibition by the clinical antivirals nirmatrelvir and ensitrelvir

**DOI:** 10.64898/2026.06.01.729201

**Authors:** Hyojin Kim, Jinsook Ahn, Juyeon Lee, Seungheon Jung, Ji Woo Kim, Byungil Kim, Nam-Chul Ha, Inseong Jo

**Affiliations:** Infectious Diseases Therapeutic Research Center, Division of Therapeutics and Biotechnology, Korea Research Institute of Chemical Technology (KRICT), Daejeon 34114, Republic of Korea; Department of Agricultural Biotechnology, and Research Institute of Agriculture and Life Sciences, CALS, Seoul National University, Seoul 08826, Republic of Korea; Department of Biological Sciences, Korea Advanced Institute for Science and Technology (KAIST), Daejeon 34141, Republic of Korea; School of Pharmacy, Sungkyunkwan University, Suwon 16419, Republic of Korea; College of Pharmacy, Chung-Ang University, Seoul 06974, Republic of Korea

**Author notes:** Corresponding authors, (NH); (IJ). Pharmus Bioscience Inc., Youngin 16890, Republic of Korea (Ji Woo Kim).

**Keywords:** HKU5-CoV, main protease, nirmatrelvir, ensitrelvir, X-ray crystallography

## Abstract

The identification of *Pipistrellus* bat coronavirus HKU5 lineage 2 (HKU5-CoV-2) as a human ACE2-adapted virus highlights the need for antiviral strategies to control emerging HKU5-related merbecoviruses. However, despite the importance of the main protease (M^pro^) as a key antiviral target, structural and biochemical characterization of HKU5-CoV M^pro^ in the context of clinical inhibitors has remained limited. Here, we obtained high-resolution crystal structures of HKU5-CoV-1 M^pro^ in its apo state (1.75 Å) and in complex with the approved covalent inhibitor nirmatrelvir (1.91 Å) and the non-covalent inhibitor ensitrelvir (1.55 Å). These structures reveal an induced-fit mechanism, in which the S2 loop containing Met49 adopts an open conformation in the apo state and undergoes inhibitor-specific conformational changes upon nirmatrelvir and ensitrelvir binding. These structures served as a foundation for the characterization of HKU5-CoV-2 M^pro^ via modeling and molecular dynamics simulations. Biochemical assays revealed that HKU5-CoV-1 and HKU5-CoV-2 M^pro^ exhibited nearly identical kinetic profiles, with turnover rates approximately two-fold higher than SARS-CoV-2 M^pro^. Structural analysis revealed a highly conserved S1 subsite but distinct local environments in the S1′, S2, and S4 substrate-binding sites relevant to inhibitor recognition. Despite these variations, nirmatrelvir and ensitrelvir showed potent inhibitory activity, with comparable double-digit nanomolar IC_50_ values across all three M^pro^ proteins. Computational analyses indicated that HKU5-CoV-2 M^pro^ engages nirmatrelvir and ensitrelvir in binding modes comparable to those observed for HKU5-CoV-1, supporting a conserved mechanism of inhibitor recognition and providing a structural basis for developing pan-coronavirus antivirals targeting emerging merbecoviruses.

## Introduction

The persistent emergence of zoonotic coronaviruses (CoVs) has introduced numerous challenges to global public health, as demonstrated by the severe outbreaks of severe acute respiratory syndrome (SARS), Middle East respiratory syndrome (MERS), and the coronavirus disease 2019 (COVID-19) pandemic [1]. Their initial transmission to human populations frequently involves specific intermediate hosts, such as civets or dromedary camels for SARS-CoV or MERS-CoV, respectively; however, extensive phylogenetic analyses have revealed that these highly pathogenic viruses are believed to ultimately originate from wildlife reservoirs, particularly bats [2-9]. Bats serve as unique ecological hosts, harboring various CoVs across the *Alpha and Betacoronavirus* genera [5]. Within the *Betacoronavirus* genus, *Merbecoviruses* are concerning owing to their demonstrated potential for cross-species transmission and high fatality rates [10, 11].

Among the recognized bat-borne *Merbecoviruses, Pipistrellus* bat CoV HKU5 (HKU5-CoV) is closely related to MERS-CoV [12, 13]. Historically, HKU5-CoV has been regarded as a bat-restricted virus with minimal threat to humans; however, the recent discovery of a distinct lineage, HKU5-CoV-2, has significantly altered this perspective. Unlike its predecessor, HKU5-CoV-2 can efficiently utilize human angiotensin-converting enzyme 2 (ACE2) as a functional entry receptor [14-16]. Combined with its robust infectivity across diverse mammalian ACE2 orthologs, this ability suggests that HKU5-related *Merbecoviruses* constitute a dynamic evolutionary interface, whereby minor genetic changes can expand their host range [15, 16]. Consequently, characterizing the molecular architecture of this lineage is important to pandemic preparedness.

The sustained human transmission potential of ACE2-adapted viruses, such as HKU5-CoV-2, highlights the necessity of a proactive strategy for assessing and repurposing CoV therapeutics that were initially developed for SARS-CoV-2 [14, 17, 18]. Moreover, establishing a robust preliminary defense is critical prior to any spillover event that could cause widespread infection [18, 19]. Owing to the high conservation of viral replication machinery, targeting essential components shared by both bat and human CoVs remains a viable strategy for pan-CoV interventions [14, 20, 21].

Among these targets, the main protease (M^pro^), also known as the 3C-like protease, is appealing because of its critical role in the viral life cycle. The M^pro^ is responsible for cleaving large viral polyproteins (pp1a and pp1ab) into functional non-structural proteins necessary for replication (Fig. 1A) [22-26]. Owing to its conserved 3C-like architecture and essential catalytic function, the M^pro^ has been extensively studied in antiviral research, enabling the identification of various covalent and non-covalent inhibitors and numerous high-resolution complex structures across diverse RNA viruses [27-35].

**Figure 1.**
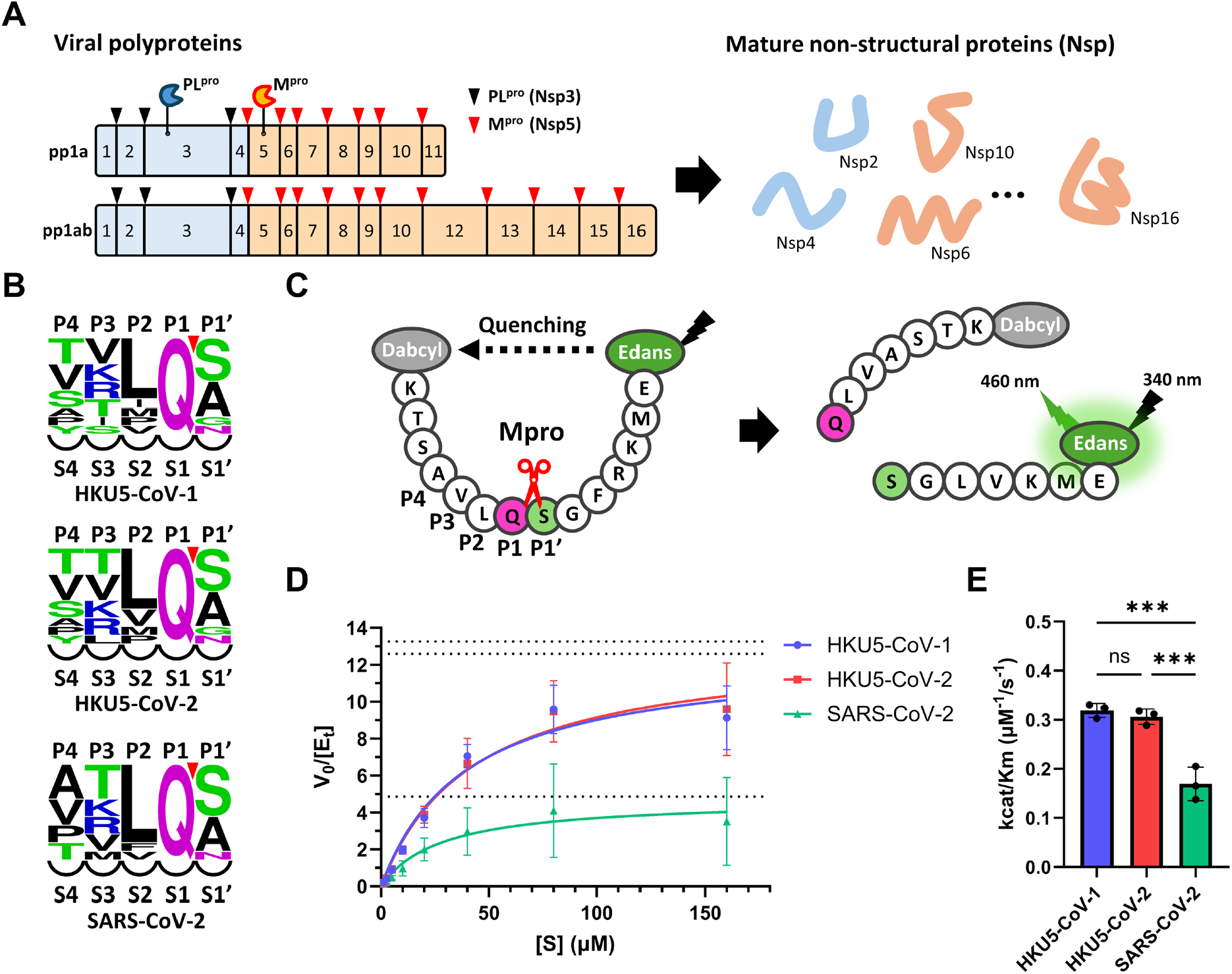
Enzymatic characterization of M^pro^ proteins. **(A)** Schematic of viral polyprotein cleavage into non-structural proteins (Nsp1-16) by PL^pro^ (black triangles) and M^pro^(red triangles) **(B)** Sequence comparisons of the autocleavage sites for M^pro^ across HKU5-CoV-1, HKU5-CoV-2, and SARS-CoV-2. The logo plots were generated by WebLogo [63] **(C)** Schematic diagram of the FRET-based enzymatic assay for the M^pro^ using a dabcyl (N-terminus)- and edans (C-terminus)-labeled peptide substrate, resulting in increased fluorescence upon proteolytic cleavage. **(D)** Michaelis–Menten curve derived from initial velocity measurements at varying substrate concentrations. Values are means ± SEM of at least three independent experiments performed in triplicates and fitted with the Michaelis–Menten equation. **(E)** Comparison of catalytic efficiencies (k_cat_/K_m_) for M^pro^ of HKU5-CoV-1 (blue), HKU5-CoV-2 (red), and SARS-CoV-2 (green). Data are presented as mean ± SD (n = 3); statistical significance was determined using one-way analysis of variance with Tukey′s multiple comparisons test (*\*P < 0*.*05, **P < 0*.*01, ***P < 0*.*001*) using GraphPad Prism 10.

The clinical efficacy of the M^pro^ inhibitors nirmatrelvir and ensitrelvir against SARS-CoV-2 has been demonstrated in clinical trials, leading to their approval as COVID-19 therapeutics [36-42]. Although nirmatrelvir has been reported to inhibit ACE2-adapted HKU5-CoV-2 at the cellular level, structural and biochemical data on the inhibition mechanism of HKU5-CoV M^pro^ remain limited [14]. Therefore, it is essential to identify which clinical M^pro^ inhibitors can effectively block HKU5-CoV main proteases and to elucidate their molecular mechanisms of inhibition through comparative structural and biochemical analyses, in order to assess the robustness of frontline protease-targeting antivirals against emerging zoonotic merbecoviruses.

Here, we report the first crystal structures of HKU5-CoV-1 Mpro in the apo form and in complex with nirmatrelvir and ensitrelvir, and integrate biochemical and computational analyses to define the molecular basis of inhibitor recognition across HKU5-CoV lineages.

## Results

### Identification of the M^pro^ from HKU5-CoV lineages 1 and 2

The M^pro^ (Nsp5) of HKU5-CoV-2 demonstrated 93% and 50% amino acid sequence identities with those of HKU5-CoV-1 and SARS-CoV-2, respectively. To elucidate the substrate specificity of these proteases, 11 canonical polyprotein cleavage junctions (Nsp4/5 to Nsp15/16) from HKU5-CoV-1, HKU5-CoV-2, and SARS-CoV-2 were aligned **(Figs. 1A and 1B, Table S1)**. The cleavage sequences at these 11 sites were highly conserved between HKU5-CoV-1 and HKU5-CoV-2, with only a limited number of lineage-specific substitutions **(Table S1)**. Across all three viruses, the P1 position consistently featured Gln at all 11 junctions, aligning with the canonical substrate specificity of CoV M^pro^ proteins. The P1′ position was predominantly occupied by small, uncharged residues (Ser/Ala/Gly), except at the Nsp8/9 junction, where the N-terminal Asn residue of Nsp9 served as the substrate for NiRAN-mediated NMPylation. At P2, the viruses consistently maintained hydrophobic residues with distinct distribution patterns. SARS-CoV-2 showed the strongest preference for Leu (9 of 11 sites), followed by Phe and Val. Conversely, HKU5-CoV-1/2 exhibited a broader hydrophobic repertoire. Although Leu was still predominant (7 of 11 sites), the remaining positions were diversified among Met, Val, Pro, and Ile, indicating a conserved hydrophobic preference but with greater sequence diversity **(Fig. 1B and Table S1)**. At P4, the SARS-CoV-2 M^pro^ restricted substrate recognition to small amino acids, including Ala, Val, Thr, and Pro, whereas the HKU5-CoV-1/2 M^pro^ proteins recognized these small residues while accommodating the bulky aromatic Tyr at the Nsp13/14 junction **(Fig. 1B and Table S1)**.

Therefore, the HKU5-CoV M^pro^ proteins retained the canonical substrate-recognition features of CoV M^pro^ proteins, particularly an invariant P1 glutamine, conserved hydrophobic preference at P2, and a small uncharged residue at P1′ across all cleavage junctions.

To assess the enzymatic characteristics of the HKU5-CoV-1/2 and SARS-CoV-2 M^pro^, steady-state kinetic parameters were determined using a fluorescence resonance energy transfer (FRET)-based peptide substrate [(Dabcyl)KTSAVLQ↓SGFRKME(Edans)] that included the conserved P2–P1–P1′ sequence (L–Q–S). The M^pro^ proteins of the two HKU5 lineages exhibited nearly identical kinetic profiles. For example, the HKU5-CoV-1 M^pro^ demonstrated a K_m_ and catalytic efficiency (k_cat_/K_m_) of 39.5 ± 4.0 μM and 0.32 μM^−1^ s^−1^, respectively, while those of the HKU5-CoV-2 M^pro^ were 43.7 ± 12.9 μM and 0.31 μM^−1^ s^−1^ **(Table 1)**. Considering the SARS-CoV-2 M^pro^ (K_m_ = 29.4 ± 11.7 μM; k_cat_/K_m_ = 0.17 μM^−1^ s^−1^), the K_m_ values were similar among the three proteases (29.4–43.7 μM; *p* > 0.05), suggesting comparable substrate affinity. However, both HKU5 M^pro^ variants exhibited significantly higher catalytic efficiencies due to increased turnover rates (k_cat_ = 12.6 s^−1^ for HKU5-CoV-1 and 13.3 s^−1^ for HKU5-CoV-2 vs. 4.9 s^−1^ for SARS-CoV-2; **Table 1 and Figs. 1D and 1E**). The catalytic efficiencies of the HKU5-CoV-1 and HKU5-CoV-2 M^pro^ proteins were approximately 1.9- and 1.8-fold higher, respectively, than that of the SARS-CoV-2 M^pro^ (*p* < 0.05; **Table 1 and Fig. 1E**). Therefore, the HKU5-CoV lineages encoded catalytically proficient M^pro^ proteins with substrate binding affinities comparable to that of the SARS-CoV-2 M^pro^, achieving approximately two-fold higher catalytic efficiency through substantially increased turnover rates.

**Table 1.** Steady-state kinetic parameters of the HKU5-CoV-1/2 and SARS-CoV-2 M^pro^ using the FRET substrate [Dabcyl-KTSAVLQ-SGFRKME-Edans].

| Protease | K<sub>m</sub><br>(μM) | V<sub>max</sub><br>(μM/s) | k<sub>cat</sub><br>(s<sup>-1</sup>) | k<sub>cat</sub>/K<sub>m</sub><br>(μM<sup>-1</sup>/s<sup>-1</sup>) | Relative<br>Efficiency |
| --- | --- | --- | --- | --- | --- |
| SARS-CoV-2 M<sup>pro</sup> | 29.4±11.7 | 0.49±0.16 | 4.9 | 0.17 | 1.0 |
| HKU5-CoV-1 M<sup>pro</sup> | 39.5±4.0 | 1.26±0.15 | 12.6 | 0.32 | 1.9 |
| HKU5-CoV-2 M<sup>pro</sup> | 43.7±12.9 | 1.33±0.32 | 13.3 | 0.31 | 1.8 |

### Overall structure of the apo-M^pro^ from HKU5-CoV

To determine the molecular structures of the HKU5-CoV M^pro^ proteins, crystallization screening was performed using recombinant constructs from both HKU5-CoV-1 and HKU5-CoV-2, which share 93% amino acid sequence identity and display similar enzymatic profiles **(Table 1 and Fig. 2A)**. Despite these similarities and extensive screening across various crystallization conditions, diffraction-quality crystals were obtained only for the HKU5-CoV-1 M^pro^. The crystal structure of the apo-state of HKU5-CoV-1 M^pro^ (PDB ID 9VVC) was determined at 1.75 Å resolution in space group C2 **(Table 2)**, and the asymmetric unit contained a single protomer with the canonical three-domain architecture conserved among CoV M^pro^ proteins **(Fig. 2B)**. Domains I (residues 10–98) and II (residues 99–199) adopted the chymotrypsin-like fold, comprising two antiparallel β-barrels, with the substrate-binding cleft and catalytic site positioned at the interdomain interface. N-finger (residues 1–9) and domain III (residues 200–306) were essential for dimerization **(Figs. 2A and 2B)**.

**Table 2.**
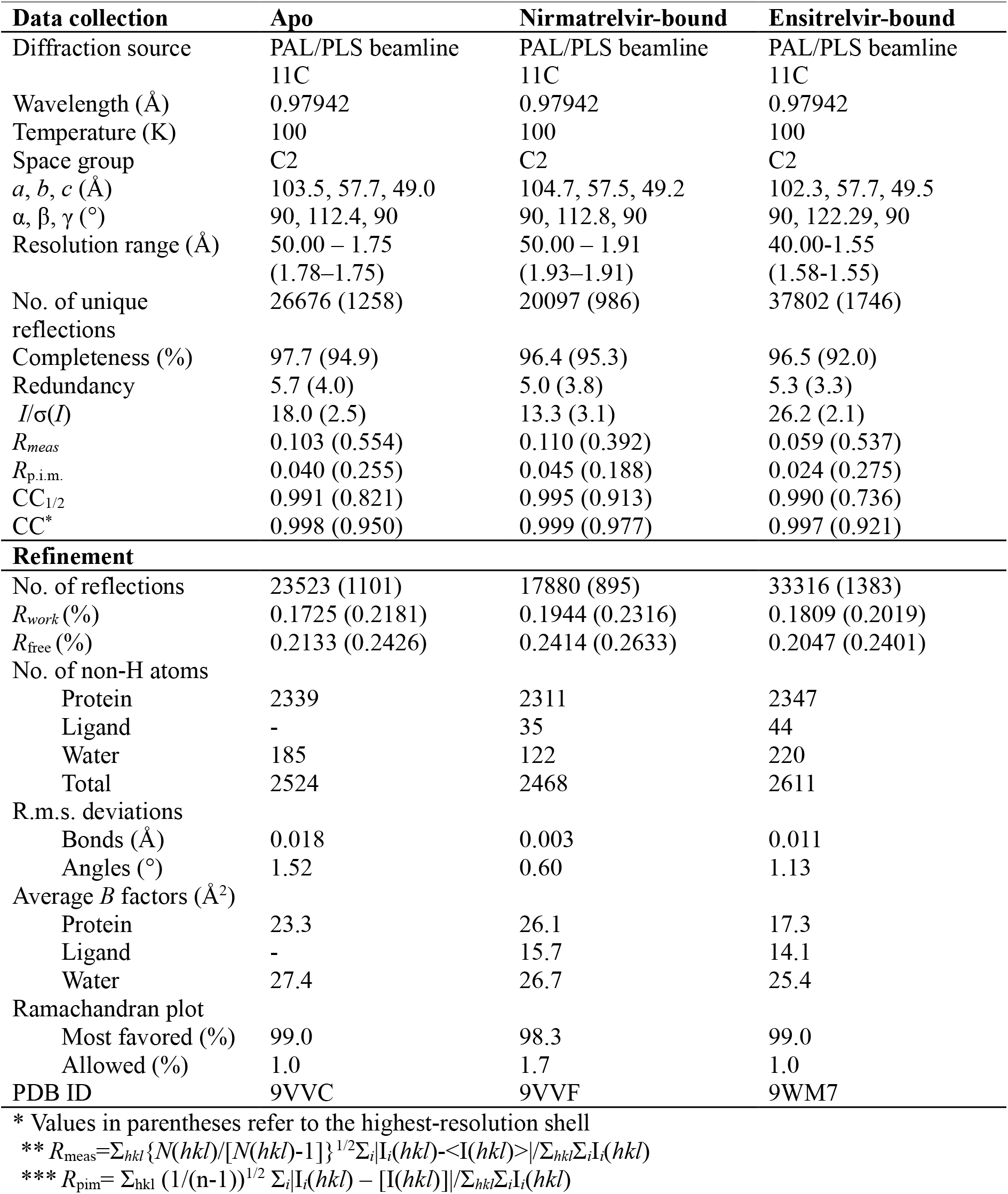
Data collection and refinement statistics for apo, nirmatrelvir-bound, and ensitrelvir-bound HKU5-CoV-1 M^pro^ structures.

| **Data collection** | **Apo** | **Nirmatrelvir-bound** | **Ensitrelvir-bound** |
| --- | --- | --- | --- |
| Diffraction source | PAL/PLS beamline<br>11C | PAL/PLS beamline<br>11C | PAL/PLS beamline<br>11C |
| Wavelength (Å) | 0.97942 | 0.97942 | 0.97942 |
| Temperature (K) | 100 | 100 | 100 |
| Space group | C2 | C2 | C2 |
| *a*, *b*, *c* (Å) | 103.5, 57.7, 49.0 | 104.7, 57.5, 49.2 | 102.3, 57.7, 49.5 |
| $\alpha$, $\beta$, $\gamma$ (°) | 90, 112.4, 90 | 90, 112.8, 90 | 90, 122.29, 90 |
| Resolution range (Å) | 50.00 – 1.75<br>(1.78–1.75) | 50.00 – 1.91<br>(1.93–1.91) | 40.00–1.55<br>(1.58–1.55) |
| No. of unique reflections | 26676 (1258) | 20097 (986) | 37802 (1746) |
| Completeness (%) | 97.7 (94.9) | 96.4 (95.3) | 96.5 (92.0) |
| Redundancy | 5.7 (4.0) | 5.0 (3.8) | 5.3 (3.3) |
| *I*/$\sigma$(*I*) | 18.0 (2.5) | 13.3 (3.1) | 26.2 (2.1) |
| *R*<sub>meas</sub> | 0.103 (0.554) | 0.110 (0.392) | 0.059 (0.537) |
| *R*<sub>p.i.m.</sub> | 0.040 (0.255) | 0.045 (0.188) | 0.024 (0.275) |
| CC<sub>1/2</sub> | 0.991 (0.821) | 0.995 (0.913) | 0.990 (0.736) |
| CC* | 0.998 (0.950) | 0.999 (0.977) | 0.997 (0.921) |
| **Refinement** |  |  |  |
| No. of reflections | 23523 (1101) | 17880 (895) | 33316 (1383) |
| *R*<sub>work</sub> (%) | 0.1725 (0.2181) | 0.1944 (0.2316) | 0.1809 (0.2019) |
| *R*<sub>free</sub> (%) | 0.2133 (0.2426) | 0.2414 (0.2633) | 0.2047 (0.2401) |
| No. of non-H atoms |  |  |  |
| Protein | 2339 | 2311 | 2347 |
| Ligand | - | 35 | 44 |
| Water | 185 | 122 | 220 |
| Total | 2524 | 2468 | 2611 |
| R.m.s. deviations |  |  |  |
| Bonds (Å) | 0.018 | 0.003 | 0.011 |
| Angles (°) | 1.52 | 0.60 | 1.13 |
| Average *B* factors (Å<sup>2</sup>) |  |  |  |
| Protein | 23.3 | 26.1 | 17.3 |
| Ligand | - | 15.7 | 14.1 |
| Water | 27.4 | 26.7 | 25.4 |
| Ramachandran plot |  |  |  |
| Most favored (%) | 99.0 | 98.3 | 99.0 |
| Allowed (%) | 1.0 | 1.7 | 1.0 |
| PDB ID | 9VVC | 9VVF | 9WM7 |

\* Values in parentheses refer to the highest-resolution shell
\*\* $R_{\text{meas}} = \sum_{hkl} \{N(hkl)/[N(hkl)-1]\}^{1/2} \sum_i |I_i(hkl) - \langle I(hkl) \rangle| / \sum_{hkl} \sum_i I_i(hkl)$
\*\*\* $R_{\text{pim}} = \sum_{hkl} (1/(n-1))^{1/2} \sum_i |I_i(hkl) - [I(hkl)]| / \sum_{hkl} \sum_i I_i(hkl)$

**Figure 2.**
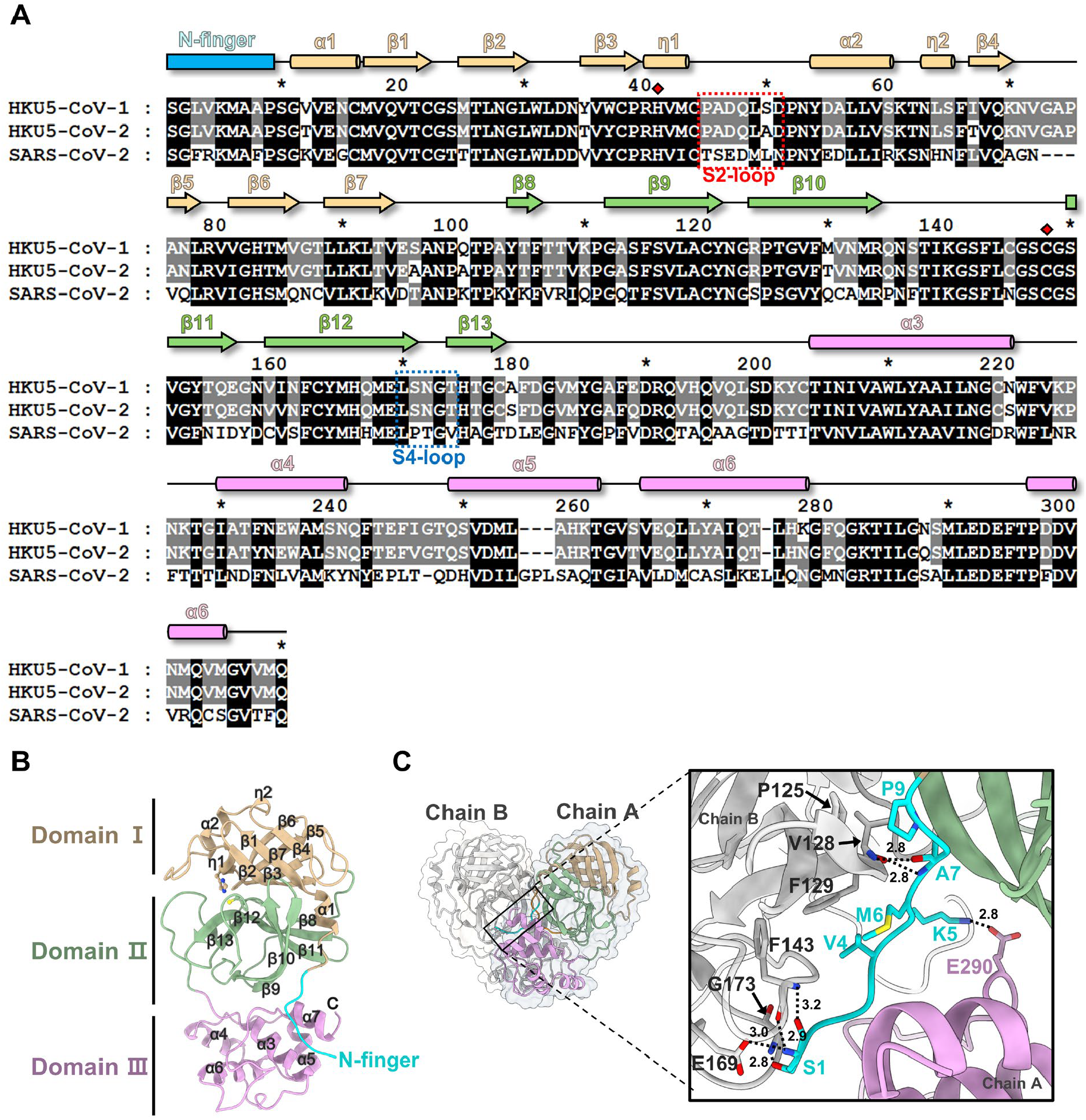
Sequence and structural features of the HKU5-CoV-1 M^pro^. **(A)** Sequence alignment of the M^pro^ enzymes from HKU5-CoV-1/2 and SARS-CoV-2. Conserved residues are shaded in gray and black, and catalytic residues are highlighted with red diamonds. S2-loop (red dotted box) and S4-loop (blue dotted box) were marked. **(B)** An asymmetric unit of the crystal structure of the HKU5-CoV-1 Apo-M^pro^. N-finger, domains I, II, and III are shown in cyan, tan, green, and pink, respectively, and the catalytic residues (His41 and Cys148) are displayed in stick representation. **(C)** Homodimer structure and interface of the HKU5-CoV-1 M^pro^. The residues involved in the dimeric interface are displayed in stick representation. The N-finger, domain II, and domain III of chain A are colored cyan, green, and pink, respectively, and chain B is shown in gray.

The biologically active homodimer formed through crystallographic symmetry and was stabilized by a cooperative network of interactions centered on the N-finger **(Figs. 2B and 2C)**. An intra-protomer salt bridge between Lys5 and Glu290 of the N-finger and domain III, respectively, anchored the N-finger. At the dimer interface, the N-terminal Ser1 formed a complex hydrogen-bonding network with the adjacent protomer. Particularly, the side chain of Ser1 interacted with the side chain of Glu169 (structurally equivalent to conserved Glu166 in the SARS-CoV-2 M^pro^) and the backbone amide of Gly173. This polar network was reinforced by reciprocal backbone–backbone interactions between Ala7 of the N-finger and Val128 of the partner protomer, ensuring precise interface alignment. Hydrophobic packing provided additional stabilization, with the N-finger engaging in hydrophobic interactions with the adjacent protomer, where Val4 and Met6 interact with Phe129, whereas Pro9 formed additional contacts with Pro125 and Val128 (**Fig. 2C**). Collectively, these multifaceted interactions generated a stable dimer interface that buried at ~1,298 Å^2^ per monomer.

### Conservation and divergence of active sites between HKU5-CoV and SARS-CoV-2 M^pro^

We used the apo structure of SARS-CoV-2 M^pro^ (PDB: 6M03), the primary clinical benchmark for current antivirals, as a reference to compare the active sites of the HKU5-CoV M^pro^ proteins [25]. Superposition of the two structures revealed a highly conserved global architecture, with individual protomers and biological dimers aligning at root-mean-square deviations (RMSDs) of 0.555 Å over 233 Cα atoms and 1.046 Å over 524 Cα atoms, respectively (**Fig. S1A**). The catalytic dyad residues, His41 and Cys148, in HKU5-CoV precisely overlapped with their SARS-CoV-2 equivalents (His41 and Cys145), revealing similar side-chain conformations. Moreover, the oxyanion hole was conserved, with the backbone amide nitrogen atoms of Gly146, Ser147, and Cys148 in HKU5-CoV (corresponding to Gly143, Ser144, and Cys145 in SARS-CoV-2) adopting superimposable positions (**Fig. S1B**).

The protease active site comprised five discrete subsites (S4–S3–S2–S1–S1′) that accommodated the corresponding P4–P3–P2–P1–P1′ residues of substrate peptides, with the catalytic dyad (Cys148–His41 in HKU5-CoVs; Cys145–His41 in SARS-CoV-2) positioned between the S1 and S1′ subsites **(Figs. 1B and 3A)**. The S1 subsite was formed by a network of conserved residues, including Phe143, His166, His175, and Glu169 from one protomer and Ser1 from the adjacent protomer **(Fig. 3B)**. The hydrogen-bonding capacity of this pocket architecture enabled the binding of two water molecules (w1 and w2), where w1 bridged the backbone carbonyl of Phe143 and Glu169, while w2 connected w1 to the His166 side chain (**Fig. 3B**). Additionally, the backbone of Phe143 and Ser1 established hydrogen bonds across the dimer interface **(Fig. 3B)**. These residues were closely superimposed with SARS-CoV-2 M^pro^ structure (structurally equivalent to Phe140, His163, His172, Glu166, and Ser1 in SARS-CoV-2), indicating extensive conservation of the S1 architecture **(Figs. 3B and F)**.

**Figure 3.**
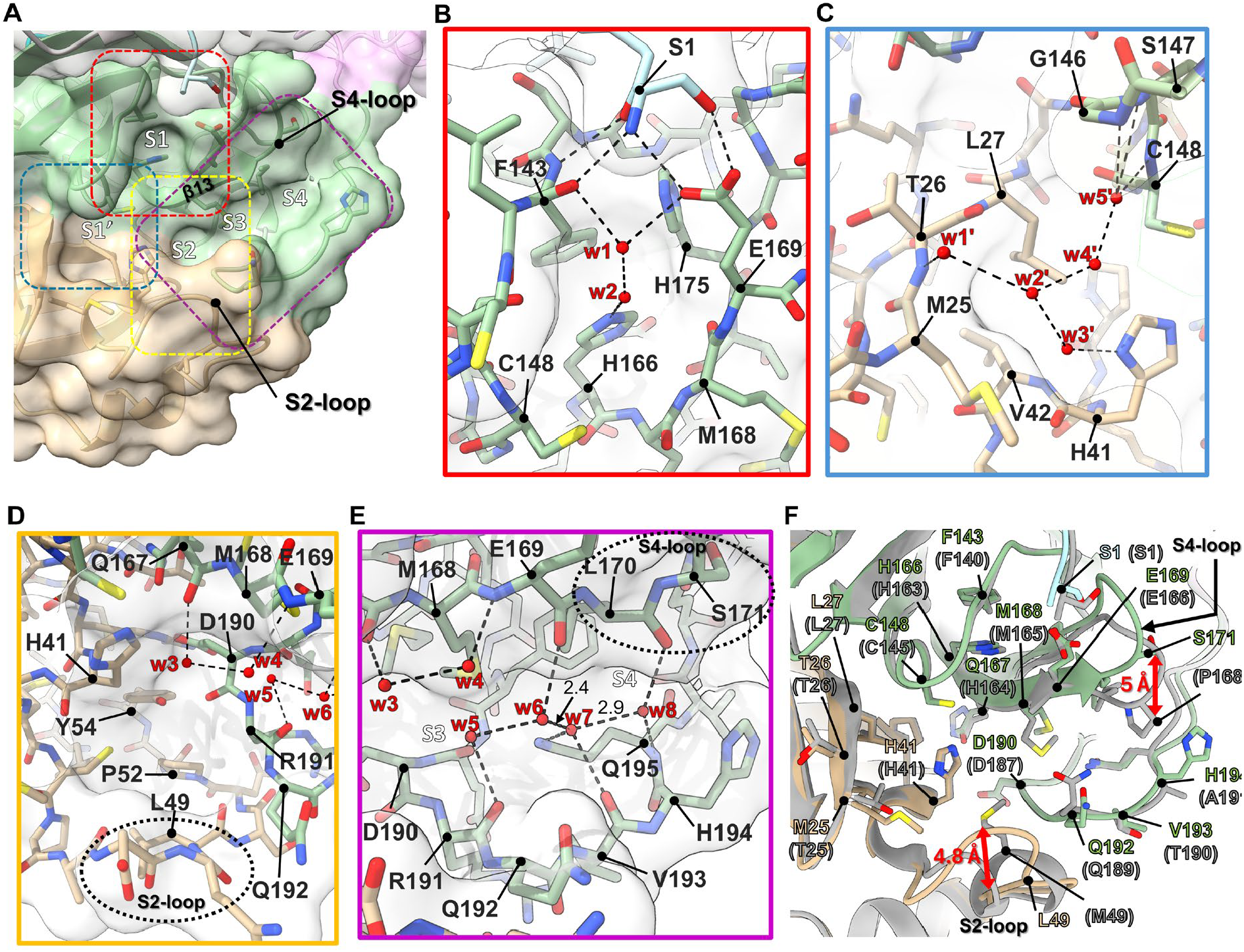
Crystal structure of the apo form of the HKU5-CoV-1 M^pro^. (**A)** Surface representation of the substrate-binding site of apo-M^pro^. The dashed boxes (red, blue, yellow, and violet) are enlarged in (B-E) panels. **(B–E)** Close-up views of the local environments of subsites S1 (B), S1′ (C), S2 (D), and S3/S4 (E), showing side chains and ordered water molecules (red spheres) engaged in hydrogen-bonding networks. Dashed lines indicate hydrogen bonds and water-mediated contacts, with distances given in Å. **(F)** Structural comparison of the active site and subsites of HKU5-CoV-1 M^pro^ (green) and SARS-CoV-2 M^pro^ (gray), representing the conserved overall architecture and subsite-defining residues, as well as subtle side-chain rearrangements and loop shifts around the S2 region.

The structural conservation of the S1 pocket is consistent with the invariant P1-glutamine specificity observed for both viruses **(Fig. 1B)**.

The S1′ subsite exhibited similar structural conservation, primarily defined by the Met25, Thr26, Leu27, His41, and Val42 residues, combined with the oxyanion-hole backbone segment comprising Cys145, Gly146, Ser147, and Cys148 (**Fig. 3C**). Particularly, five ordered water molecules (w1′–w5′) established an extensive hydrogen-bond network that connected the oxyanion hole to the backbone amide and side chain of Thr26 and His41, respectively **(Fig. 3C)**. Despite minor sequence differences in the residues, the three-dimensional organization of the hydrogen-bond donors/acceptors that coordinated this water-mediated network remained conserved between the HKU5-CoVs and SARS-CoV-2 (**Figs. 3C and 3F**).

Conversely, the S2 subsite exhibited structural differences between the M^pro^ proteins of HKU5-CoVs and SARS-CoV-2. The S2 pocket in the HKU5-CoV M^pro^ proteins comprised His41, Leu49, Pro52, Tyr54, Met168, Asp190, Arg191, and Gln192, which structurally correspond to His41, Met49, Pro52, Tyr54, Met165, Asp187, Arg188, and Gln189 in that of SARS-CoV-2 **(Figs. 3D and 3F)**. This pocket accommodated selective hydration, with w3 and w4 binding to the backbone atoms of Met168 and Glu169, whereas the deep hydrophobic region around Leu49 remained unhydrated (**Fig. 3D**). This conformational difference was supported by elevated B-factors of the S2-loop in the HKU5-CoV apo-M^pro^, indicating conformational flexibility (**Fig. S2A**). In the apo state, this loop adopted a different conformation in HKU5-CoVs, with Leu49 displaced outward by ~4.8 Å relative to Met49 in SARS-CoV-2 (measured between Cγ atoms), resulting in a more open S2 cavity (**Fig. 3E**).

Moreover, the S3–S4 region exhibited measurable divergence; however, its overall scaffold remained comparable. The S3–S4 pocket in the HKU5-CoV M^pro^ proteins comprised residues at the end of β12 (Glu169, Leu170, and Ser171) and a loop region (residues 189–195) between β13 and α3, which structurally correspond to Glu166, Leu167, Pro168, and residues 186–192 in that of SARS-CoV-2 **(Figs. 2A, 3E, and 3F)**. Within this structural framework, an extensive hydrogen-bonding network was formed by four water molecules (w5–w8). Particularly, w5 and w6 interacted with the backbone carbonyls of Arg191 and Glu169, respectively, whereas w7 interacted with the backbone carbonyl of Val193 and side chain of Gln195 and w8 interacted with the backbone carbonyl of Leu170 and backbone amide of Gln195 (**Fig. 3E)**. Despite this overall structural correspondence, the S3–S4 pocket displayed distinct structural variations resulting from structural differences in a loop region spanning residues 170–174 (referred to as the S4-loop; equivalent to SARS-CoV-2 residues 167–171). Within this loop, Ser171 in HKU5-CoV M^pro^ corresponds to Pro168 in SARS-CoV-2 M^pro^. The Ser171 residue was displaced by ~5 Å relative to Pro168 (measured between C_β_ atoms), resulting in a more expanded S4 cavity in HKU5-CoV M^pro^ than in the SARS-CoV-2 M^pro^ (**Figs. 3E and 3F**).

### Cross-species inhibitory potency and structure determination using clinical M^pro^ inhibitors

To evaluate the cross-species efficacy of clinically approved SARS-CoV-2 M^pro^ inhibitors, the inhibitory potency (IC_50_) of nirmatrelvir and ensitrelvir against all three M^pro^ proteins was determined using identical FRET-based assay conditions. Nirmatrelvir showed comparable inhibitory potency across all three enzymes, with IC_50_ values of 33.8 ± 4.2, 34.8 ± 4.8, and 39.8 ± 7.2 nM for the M^pro^ proteins of HKU5-CoV-1, HKU5-CoV-2, and SARS-CoV-2, respectively. No significant differences were detected between the HKU5 and SARS-CoV-2 M^pro^ proteins in pairwise comparisons **(Fig. 4A)**. Ensitrelvir also inhibited all three M^pro^ enzymes, with IC_50_ values of 50.3 ± 2.4, 76.2 ± 26.8, and 37.1 ± 2.9 nM for HKU5-CoV-1, HKU5-CoV-2, and SARS-CoV-2, respectively. Although the IC_50_ values for the HKU5 M^pro^ proteins tended to be higher than those for the SARS-CoV-2 M^pro^, these differences were not significant, and all values remained within the double-digit nanomolar range **(Fig. 4B)**.

**Figure 4.**
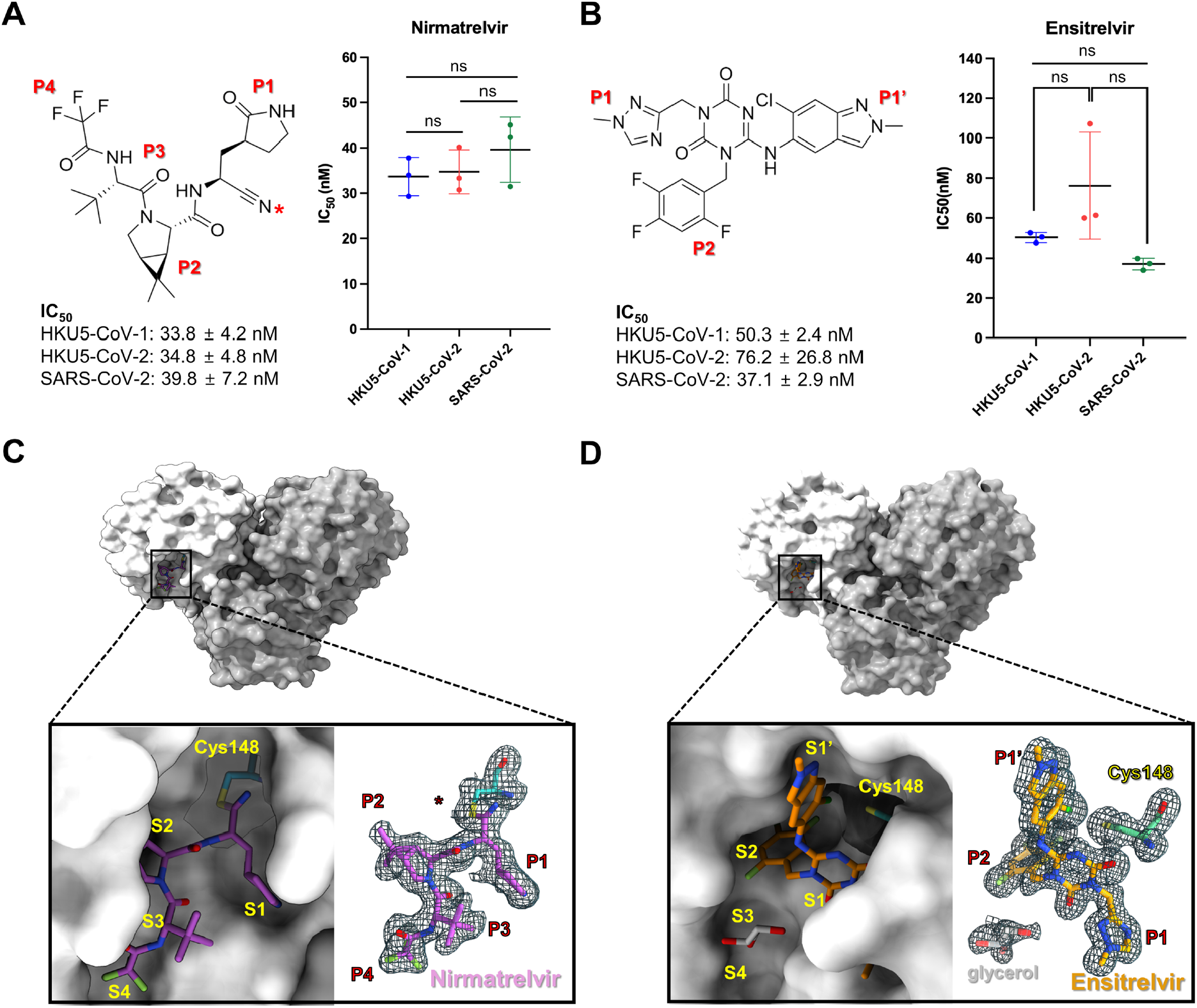
Inhibition potency and crystal structure of HKU5-CoV-1 M^pro^ in complex with nirmatrelvir and ensitrelvir. (A-B) Chemical structures of nirmatrelvir (A) and ensitrelvir (B), together with their inhibitory activities against HKU5-CoV-1, HKU5-CoV-2, and SARS-CoV-2 M^pro^. IC_50_ values were determined by FRET-based enzymatic assays and are shown as mean ± SD (n = 3). (C, D) Overall and close-up views of the crystal structures of HKU5-CoV-1 M^pro^ bound to nirmatrelvir (C) and ensitrelvir (D), showing the ligands occupying subsites. For each complex, the right panels display polder omit maps contoured at 1.0σ around the inhibitor and Cys148 (omitted from map calculation) to validate ligand placement; in the ensitrelvir complex, additional electron density corresponding to a bound glycerol molecule is also observed.

To elucidate the structural basis for these inhibition profiles, the co-crystal structures of the HKU5-CoV-1 M^pro^ in complex with nirmatrelvir (1.91 Å resolution) and ensitrelvir (1.55 Å resolution) were determined (**Table 2**). Both complexes crystallized in space group C2 with unit-cell parameters similar to the apo form (**Figs. 4C and 4D, Table 2**). In the nirmatrelvir complex, continuous electron density between Cys148 and the nitrile warhead provided direct crystallographic evidence for thioimidate bond formation, and the inhibitor extended across the S1–S4 substrate-binding subsites (**Fig. 4C**). Conversely, the ensitrelvir complex showed separated densities for the inhibitor and Cys148, consistent with non-covalent binding. Ensitrelvir primarily occupied the S1′, S1, and S2 subsites, while an ordered glycerol molecule from the crystallization solution occupied the S3–S4 region (**Fig. 4D**).

### Structural basis of nirmatrelvir recognition by the HKU5-CoV-1 M^pro^

The crystal structure of the HKU5-CoV-1 M^pro^–nirmatrelvir complex (PDB ID 9VVF) revealed covalent binding, in which the nitrile warhead formed a thioimidate adduct with catalytic Cys148 **(Fig. 5A)**. The resulting thioimidate nitrogen was positioned within the oxyanion hole formed by the backbone amides of Gly146, Ser147, and Cys148. An ordered water molecule (w1) also bridged the backbone of Gly146 to the thioimidate nitrogen. The P1 γ-lactam carbonyl of nirmatrelvir interacted with the His166 side chain, whereas the lactam amide engaged the Glu169 side chain **(Fig. 5B)**. The P1 γ-lactam-mediated interactions corresponded to the hydrogen-bonding network occupied by two ordered water molecules in the S1 site of the apo structure **(Figs. 3B and 5B)**. These P1 γ-lactam interactions with His166 and Glu169 were also conserved in the SARS-CoV-2 M^pro^–nirmatrelvir complex **(Fig. 5E)** [43]. The amide nitrogen between the P1 and P2 positions of nirmatrelvir formed a hydrogen bond with the backbone carbonyl of Gln167, and the carbonyl between the P2 and P3 positions engaged the backbone amide of Glu169, displacing the ordered water molecules that occupied these sites in the apo structure **(Figs. 3D and 5C)**. The P2 bicycloproline moiety was buried in the S2 subsite through hydrophobic contacts with Leu49, His41, Met168, and Gln192, forming a compact non-polar pocket **(Fig. 5C)**. Compared to the apo structure, nirmatrelvir binding induced a localized rearrangement of the S2-loop (residue 45– 51), shifting Leu49 by ~3.6 Å toward the P2 group to a position that closely superimposed on Met49 in the SARS-CoV-2 M^pro^–nirmatrelvir complex, resulting in a similar S2 geometry **(Figs. 5D and 5E)**.

**Figure 5.**
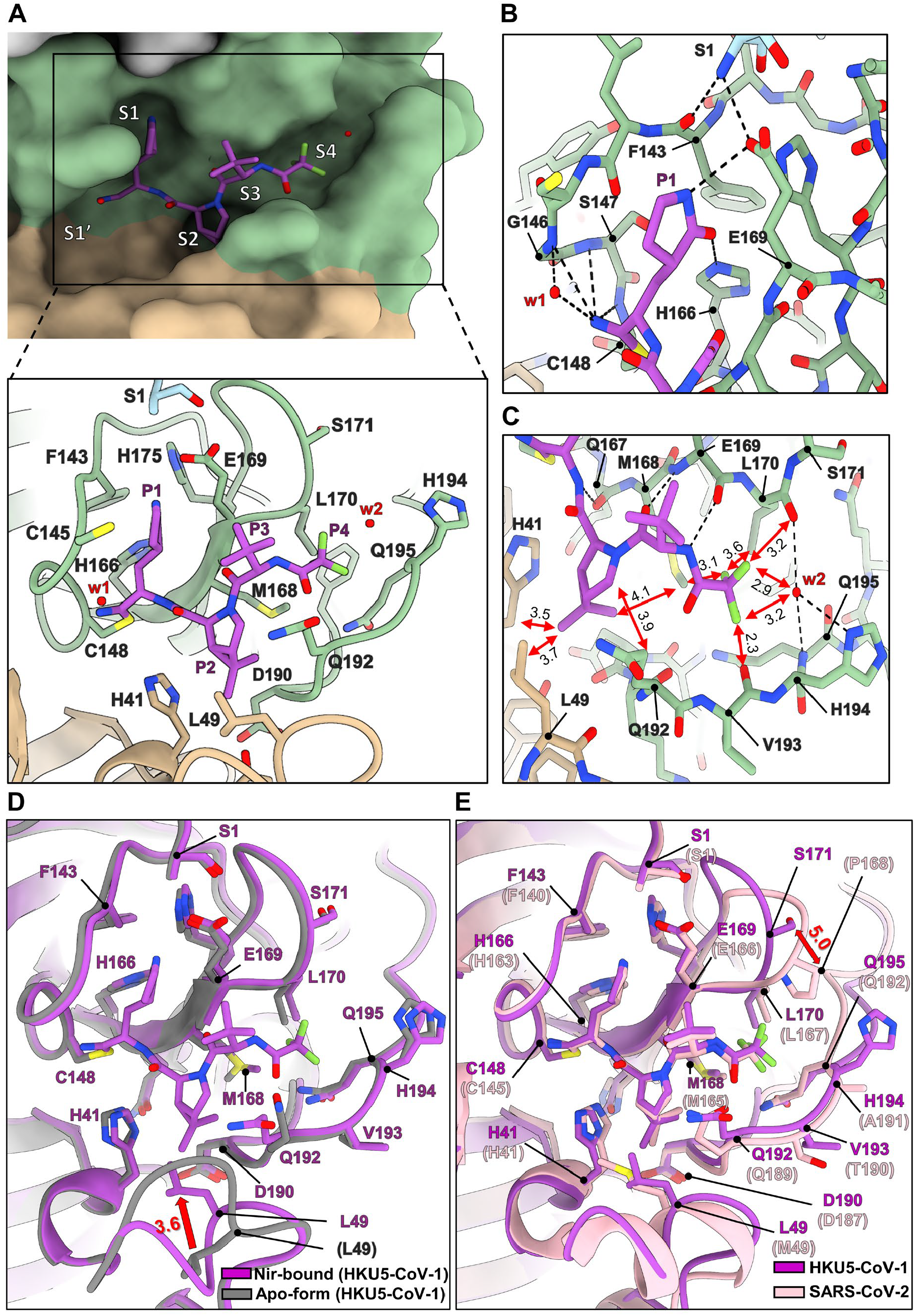
Crystal structure of the HKU5-CoV-1 M^pro^–nirmatrelvir complex. **(A)** Active site of the HKU5-CoV-1 M^pro^–nirmatrelvir complex shown as a surface representation (top) and as a ribbon representation (bottom), highlighting key active-site residues (sticks) together with nirmatrelvir (purple stick). Domains I and II are colored tan and green, respectively. **(B, C)** Close-up views of the P1 (B) and P2–P4 (C) moieties of nirmatrelvir bound in the S1 and S2–S4 subsites, respectively. Dashed lines indicate hydrogen bonds and water-mediated contacts. Red spheres represent water molecules (w1, w2). **(D)** Structural superposition of the apo-form (grey) and nirmatrelvir-bound (purple) HKU5-CoV-1 M^pro^. Residue labels are colored according to the respective structure (purple, nirmatrelvir-bound; grey, apo); equivalent residues in the apo structure are indicated in parentheses. **(E)** Structural comparison of the HKU5-CoV-1 M^pro^–nirmatrelvir complex (purple) with the SARS-CoV-2 M^pro^–nirmatrelvir complex (pink; PDB: 8DZ2). Residue labels follow the color scheme of the corresponding structure; equivalent SARS-CoV-2 M^pro^ residues are shown in parentheses.

In the S3–S4 region, the backbone of nirmatrelvir formed an additional hydrogen bond with the amide nitrogen between the P3 and P4 positions, which formed hydrogen bonds with the backbone carbonyl of Glu169 **(Fig. 5C)**. In the apo structure, this carbonyl instead formed hydrogen bonds with a water molecule **(Fig. 3E)**. The P4 CF_3_ group occupied a cleft defined by Met168, Leu170, and Val193. One fluorine atom was located 3.1 Å from the Cε atom of Met168 and 3.6 Å from the Cδ atom of Leu170, establishing hydrophobic contacts. A second fluorine atom was positioned 2.3 Å from the Val193 backbone carbonyl oxygen and 3.2 Å from an ordered water molecule (w2) that was coordinated by the Leu170 backbone carbonyl, Gln192 backbone amide, and His194 side chain. The third fluorine atom was located 3.2 and 2.9 Å from the Leu170 backbone carbonyl and w2, respectively **(Fig. 5C)**.

A comparison with the apo HKU5-CoV-1 M^pro^ structure revealed no significant conformational changes in the S3–S4 region upon nirmatrelvir binding **(Fig. 5D)**. Conversely, superposition with the SARS-CoV-2 M^pro^–nirmatrelvir complex revealed subtle differences arising from a divergence in the S4-loop **(Fig. 5E)**. Similarly, Pro168 in the apo SARS-CoV-2 M^pro^ (equivalent to Ser171 in HKU5-CoV-1) contributed to the narrow S4 cavity, precluding water coordination at this position **(Figs. 5C and 5E)**. Consequently, the fluorine atoms (CF_3_) in the SARS-CoV-2 complex were primarily surrounded by hydrophobic atoms, whereas in HKU5-CoV-1, they were located in a relatively more polar environment, consistent with their proximity to backbone carbonyl and ordered water.

### Structural basis of ensitrelvir binding to the HKU5-CoV-1 M^pro^

The crystal structure of the ensitrelvir-bound HKU5-CoV-1 M^pro^ (PDB ID 9WM7) revealed non-covalent binding by the inhibitor, with its core 1,3,5-triazine-2,4-dione ring serving as a scaffold that positioned the 6-chloro-2-methyl-indazole (P1′), 1-methyl-1,2,4-triazole (P1), and 2,4,5-trifluorophenyl (P2) moieties into the S1′, S1, and S2 subsites, respectively **(Figs. 4B and 6A)**. The central heterocycle moiety was positioned such that one of its carbonyl oxygens occupied the oxyanion hole formed by the backbone amides of Gly146, Ser147, and Cys148 (3.0–3.2 Å). A second core carbonyl formed a hydrogen bond with the backbone amide of Glu169 (3.3 Å), and an ordered water molecule (w1) bridged the backbone amide of Gly146 to a ring nitrogen of the core scaffold, providing an additional water-mediated contact **(Fig. 6B)**. The P1 triazole interacted with the His166 side chain, recapitulating the S1 recognition pattern characteristic of nirmatrelvir, whereas no direct contact was observed with the Glu169 side chain **(Figs 5B and 6B)**. Compared with the nirmatrelvir-bound HKU5-CoV-1 M^pro^, engagement of the oxyanion hole and Glu169 by the core carbonyls was accompanied by only subtle rearrangements of the Gly146–Ser147– Cys148 segment and β12 strand, displacing the backbone amides of Gly146 and Glu169 by ~0.5–0.6 Å, whereas the remaining S1 pocket residues remained largely unperturbed **(Fig. 6D)**.

**Figure 6.**
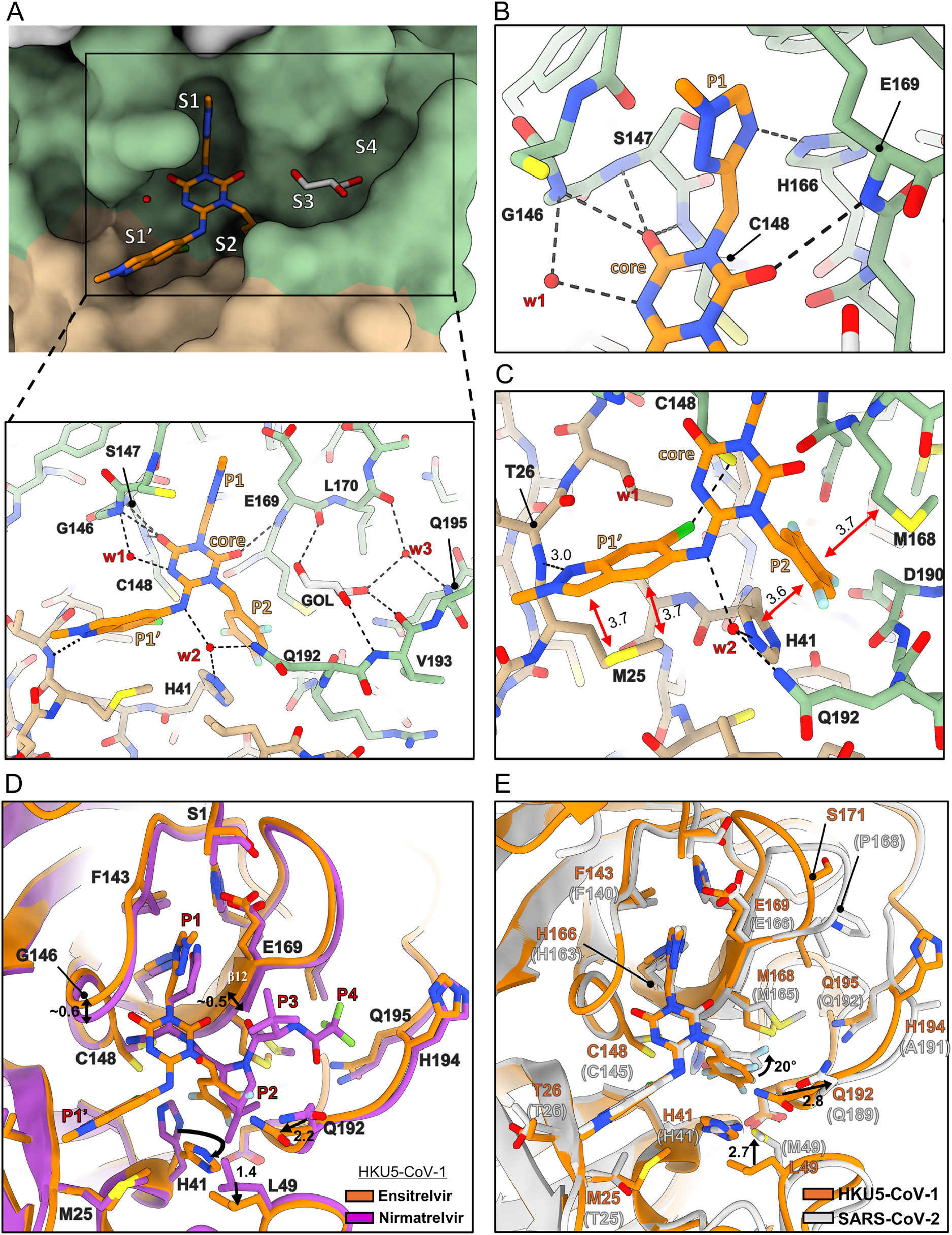
Crystal structure of the HKU5-CoV-1 M^pro^–ensitrelvir complex. **(A)** Active site of the HKU5-CoV-1 M^pro^–ensitrelvir complex shown as a surface representation (top) and as a ribbon representation (bottom), highlighting key active-site residues together with ensitrelvir (orange sticks) and a glycerol molecule (gray sticks). Domains I and II are colored tan and green, respectively. **(B, C**) Close-up views of the P1 moiety (B) and the P1′ and P2 moieties (C) of ensitrelvir bound in the S1, S1′, and S2 subsites, respectively. Dashed lines indicate hydrogen bonds. Red spheres denote water molecules (w1–w3), and glycerol (GOL) is shown as gray sticks. **(D)** Structural comparison of the HKU5-CoV-1 M^pro^ complexes with ensitrelvir (orange) and nirmatrelvir (purple). The P1–P4 and P1′ substituents of the inhibitors are labeled, and residue labels are colored according to the corresponding complex (orange, ensitrelvir-bound; purple, nirmatrelvir-bound). Conserved active-site water molecules (w1–w3) are shown as red spheres. **(E)** Structural comparison of the HKU5-CoV-1 M^pro^–ensitrelvir complex (orange) with the SARS-CoV-2 M^pro^–ensitrelvir complex (gray; PDB ID:8DZ0). Residue labels follow the color scheme of the corresponding structure; equivalent residues in SARS-CoV-2 M^pro^ are indicated in parentheses.

The P1′ 6-chloro-2-methyl-indazole group formed a hydrogen bond with the backbone amide of Thr26 (3.0 Å). The side chain of Met25 was positioned across the indazole scaffold, with its sulfur positioned over the five-membered ring and its terminal methyl group oriented toward the six-membered ring. This orientation supported simultaneous S–π and CH–π contacts anchoring the P1′ group. The 6-chloro substituent formed a halogen bond with the sulfur of Cys148 at a distance of 3.4 Å. Moreover, the linker nitrogen connecting the core and P1′ rings participated in a water-mediated hydrogen-bond network, in which an ordered water molecule (w2) bridged this nitrogen to the side chains of His41 and Gln192 **(Fig. 6C)**.

The P2 2,4,5-trifluorophenyl group occupied the S2 pocket formed by His41, Met168, and Gln192. The imidazole ring of His41 interacted with the P2 aromatic ring in a face-to-face π–π stacking arrangement, whereas the γ-carbon of Met168 engaged in a π–alkyl interaction with the trifluorophenyl ring on the opposite face of the P2 ring **(Fig. 6C)**.

Relative to the nirmatrelvir-bound HKU5-CoV-1 M^pro^, the S2 pocket adopted a remodeled geometry. For example, His41 rotated inward toward the S2 cavity, while Leu49 was displaced by ~1.4 Å outward (Cγ), and Gln192 shifted by ~2.2 Å (Cγ) toward the P2 ring **(Fig. 6D)**. In this configuration, Leu49 no longer cradled the P2 substituent, functioning instead as a structural brace between His41 and Gln192 **(Figs. 6C and 6D)**.

A comparison with the SARS-CoV-2 M^pro^–ensitrelvir complex confirmed that the inward rotation of His41, to engage in π–π stacking with the P2 aromatic ring, and the orientation of the trifluorophenyl fluorine atoms were broadly conserved between the two enzymes. However, in the SARS-CoV-2 M^pro^, Met49 (corresponding to Leu49 in the HKU5-CoV-1 M^pro^) extended 2.7 Å closer toward the P2 moiety, whereas Gln189 (equivalent to Gln192 in the HKU5-CoV-1 M^pro^) was moved 2.8 Å away from the P2 ring. Moreover, the P2 aromatic ring adopted an orientation that was rotated by ~20° relative to its pose in the HKU5-CoV-1 M^pro^. These geometric differences around S2 correlated with the altered placement of the P2 substituent. However, the S1′ interactions were largely preserved, although Thr25 replaced Met25 of the HKU5-CoV-1 M^pro^ **(Fig. 6E)**.

### Modeling studies of the HKU5-CoV-2 M^pro^ complexed with inhibitors

To characterize the binding mode of nirmatrelvir in the HKU5-CoV-2 M^pro^, an M^pro^– nirmatrelvir complex model was generated using AlphaFold3 [44]. The predicted dimer displayed high pLDDT scores and low PAE values, indicating that the overall fold and active-site geometry were reliably predicted **(Figs. S3A and B)**. When the HKU5-CoV-2 M^pro^– nirmatrelvir model was superposed onto the HKU5-CoV-1 M^pro^–nirmatrelvir crystal structure, the binding pose was closely conserved, with the nitrile warhead directed toward Cys148 and the P1–P4 substituents occupying the S1–S4 subsites in a similar arrangement **(Fig. S4A)**.

To obtain a dynamically equilibrated model, the AlphaFold3-derived HKU5-CoV-2 M^pro^–nirmatrelvir complex was subjected to explicit-solvent MD simulations using GROMACS for 50 ns. After 20 ns, both the protein backbone RMSD and that of nirmatrelvir rapidly converged and remained <0.2 nm, consistent with an energetically stable binding mode **(Fig. S4B)**. To extract a representative conformation from the equilibrated ensemble, the 20–50 ns trajectory was clustered based on the RMSD (cutoff of 0.13 nm) of active-site residues and the bound nirmatrelvir. This analysis partitioned 300 structures into four clusters. The largest cluster contained 276 frames, indicating a dominant and well-defined binding conformation, and its centroid was selected as the representative HKU5-CoV-2 M^pro^– nirmatrelvir model **(Fig. 7A)**.

**Figure 7.**
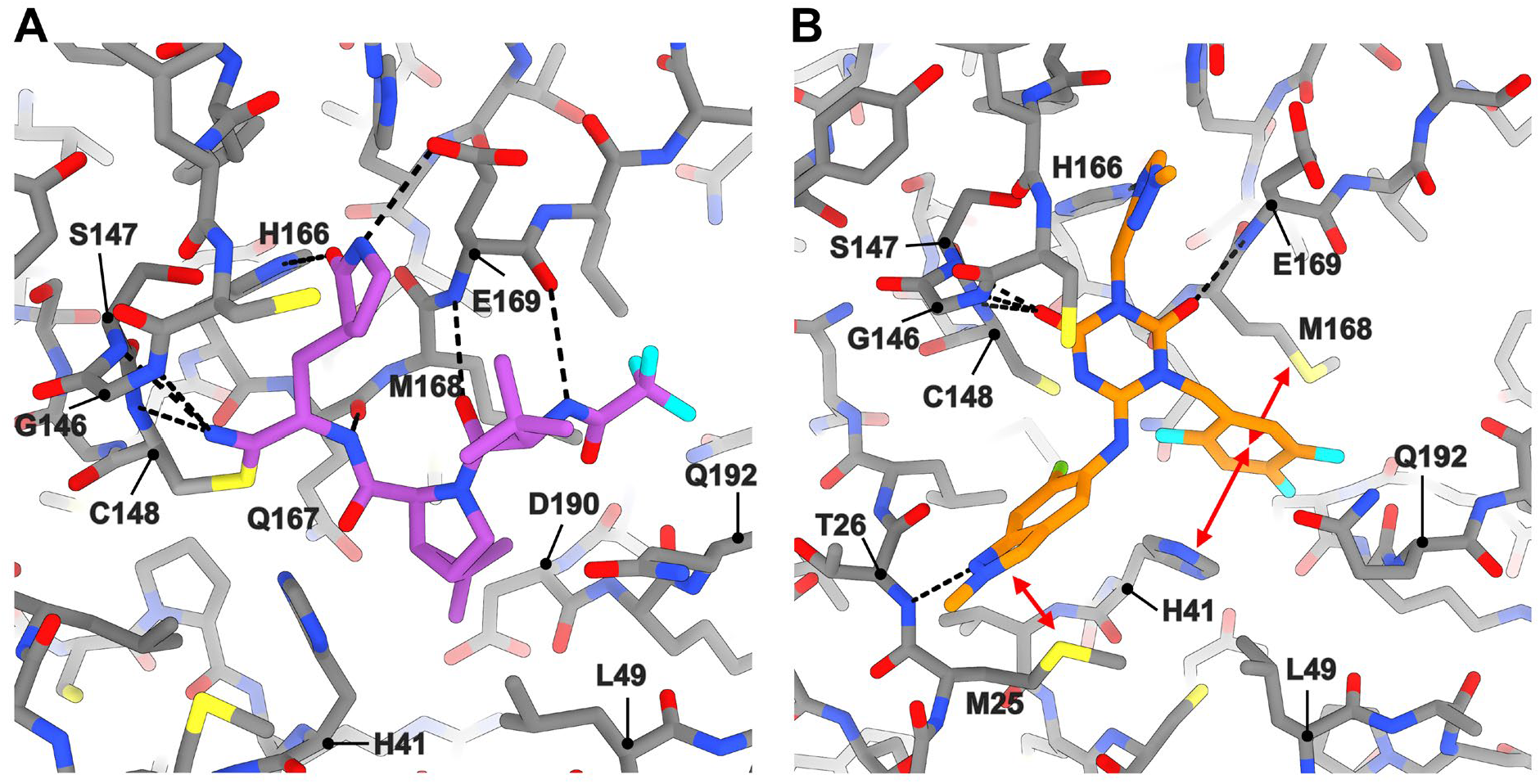
MD simulation-derived models of HKU5-CoV-2 M^pro^ complexed with nirmatrelvir and ensitrelvir. **(A)** Representative conformation of the HKU5-CoV-2 M^pro^–nirmatrelvir complex extracted from the dominant cluster of a 50-ns MD simulation trajectory. Nirmatrelvir is shown as purple sticks and key active-site residues as grey sticks. Dashed lines indicate hydrogen bonds. **(B)** Representative conformation of the HKU5-CoV-2 M^pro^–ensitrelvir complex derived from a crystal structure-guided model (threaded on the HKU5-CoV-1 M^pro^–ensitrelvir crystal structure) and equilibrated by a 50 ns MD simulation. Ensitrelvir is shown as orange sticks and key active-site residues as grey sticks. Dashed lines indicate hydrogen bonds.

In this structure, the covalent thioimidate linkage between Cys148 and the nitrile warhead was preserved, with the thioimidate nitrogen positioned in the oxyanion hole formed by the backbone amides of Gly146, Ser147, and Cys148. The P1 γ-lactam ring engaged the S1 subsite through hydrogen bonds with His166 and Glu169, while the P2 bicycloproline group remained buried in the hydrophobic S2 pocket lined by His41, Leu49, Met168, and Gln192. The backbone amide and carbonyl of the P3 segment formed reciprocal hydrogen bonds with those of Glu169 **(Fig. 7A)**. Thus, the overall binding mode of nirmatrelvir in the HKU5-CoV-2 M^pro^ was highly similar to that observed for HKU5-CoV-1.

For ensitrelvir, an initial HKU5-CoV-2 M^pro^–ensitrelvir complex was generated using AlphaFold3, yielding high-confidence pLDDT and PAE values **(Figs. S3C and D)**. Although the overall fold closely matched the HKU5-CoV-1 M^pro^–ensitrelvir crystal structure, only the P1 triazole moiety of ensitrelvir overlapped. Moreover, the P1′ indazole and P2 trifluorophenyl groups were markedly displaced, and the inward rotation of His41 toward the S2 pocket, which is characteristic of ensitrelvir-bound crystal structures, was not observed in the AlphaFold3 model **(Fig. S5A)**. During a 100-ns MD simulation, the AlphaFold3-based HKU5-CoV-2 M^pro^–ensitrelvir complex showed only partial relaxation toward the HKU5-CoV-1 binding mode during the MD simulation **(Fig. S5B)**. In the initial model at 0 ns, the P1′ 6-chloro-2-methyl-indazole was located outside the pocket formed by Met25 and Thr26, and its chloro substituent pointed outward. After 35 ns, the P1′ group rearranged into a pose more similar to those of the HKU5-CoV-1 and SARS-CoV-2 M^pro^–ensitrelvir complexes, establishing π-stacking with Met25 and a hydrogen bond between the P1′ nitrogen and Thr26 backbone amide. However, the P2 moiety did not establish close contacts with His41, as observed in the HKU5-CoV-1 crystal structure, even after 100 ns **(Fig. S5C)**. Although the MD trajectories indicated a progressive shift from the initial AlphaFold3 pose toward the HKU5-CoV-1 binding mode, the simulated timescale was insufficient to reach a fully equilibrated state.

Given the limited accuracy of the AlphaFold3-derived ensitrelvir pose, we instead constructed a crystal structure-guided model of the HKU5-CoV-2 M^pro^–ensitrelvir complex by threading the HKU5-CoV-2 sequence onto the HKU5-CoV-1 M^pro^–ensitrelvir crystal structure, taking advantage of the 93% sequence identity between the two proteases **(Fig. S6A)**. When this crystal structure-guided complex was subjected to a 50 ns MD simulation, both the protein backbone and bound ensitrelvir rapidly converged and remained stable, with RMSD values below 0.2 nm after ~30 ns **(Fig. S6B)**. We then extracted a representative conformation from the equilibrated ensemble by clustering the 30–50 ns segment of the trajectory on the RMSDs of the active-site residues and ensitrelvir. Using an RMSD cutoff of 0.13 nm, this procedure partitioned 200 snapshots into four clusters, and 143 frames grouped into a single dominant cluster, whose centroid was taken as the HKU5-CoV-2 M^pro^–ensitrelvir model **(Fig. 7B)**. In this structure, the P1 1-methyl-1,2,4-triazole moiety formed a conserved hydrogen bond with His166, and the carbonyl groups of the triazine–dione core scaffold interacted with the backbone amides of the M^pro^. The P1′ 6-chloro-2-methyl-indazole stacked against Met25 while donating a hydrogen bond to the Thr26 backbone amide, whose chloro group was oriented toward the catalytic Cys148 residue. The P2 2,4,5-trifluorophenyl group was deeply buried in the hydrophobic S2 pocket and engaged in tight π-stacking with the inward-rotated His41 side chain **(Fig. 7B)**. Thus, the HKU5-CoV-2 M^pro^ adopted inhibitor-binding modes that were conserved in the HKU5-CoV-1 M^pro^, providing a structurally validated framework to understand the largely similar enzymatic activity and inhibitor susceptibility of the two lineages.

## Discussion

Here, we present the first high-resolution crystal structures of HKU5-CoV-1 M^pro^ in the apo state and in complex with the clinically approved inhibitors nirmatrelvir (Paxlovid) and ensitrelvir (Xocova). Previous cellular studies have shown that the covalent M^pro^ inhibitors nirmatrelvir and GC376 retain potent activity against HKU5-CoV-2, with EC_50_ values of 30 nM and 130 nM, respectively, for inhibition of viral replication in human cells [14]. In our enzymatic assays, nirmatrelvir inhibited HKU5-CoV-1, HKU5-CoV-2, and SARS-CoV-2 M^pro^s with IC_50_ values of roughly 30–40 nM for each enzyme, in line with its cellular potency against HKU5-CoV-2. Ensitrelvir, which had not previously been evaluated against HKU5-CoV-2, inhibited HKU5-CoV-1/2 and SARS-CoV-2 M^pro^s with IC_50_ values ranging from 40 to 80 nM. Combined with enzymatic analysis, structural comparisons, and in silico studies of HKU5-CoV-2 M^pro^, these results demonstrate that HKU5-CoVs M^pro^ proteins remain susceptible to both nirmatrelvir and ensitrelvir despite local sequence divergence.

Structurally, comparison with the SARS-CoV-2 M^pro^ showed that the active site of HKU5-CoV M^pro^ combines a conserved catalytic core with more variable peripheral pockets. The catalytic dyad, oxyanion hole, and S1 pocket are effectively superimposable on those of SARS-CoV-2 Mpro. By contrast, the S2 and S4 subsites display clear species-specific differences. In the apo structure of HKU5-CoV-1 M^pro^, the S2 loop adopted a markedly open conformation, with Leu49 rotated outward from the substrate-binding cleft. Upon the binding of nirmatrelvir or ensitrelvir, this loop undergoes an induced-fit rearrangement, resulting in the repositioning of Leu49 to establish hydrophobic contacts with P2 moiety. Consistent with this behavior, the Leu49-containing S2 loop exhibits elevated B-factors in all HKU5-CoV-1 crystal structures, including the apo and inhibitor-bound forms, indicating that S2 remains intrinsically flexible even when stabilized by ligand binding **(Fig. S2)**. A similar behavior has been observed for the Met49-containing short 3_10_ helix in SARS-CoV-2 M^pro^, where multi-temperature crystallography revealed an atypical, temperature-dependent backbone variability at this region, suggesting that the Leu49/Met49 loop is capable of sampling multiple conformational states in coronavirus main proteases [45]. The S4 subsite was expanded in HKU5-CoV M^pro^ owing to the Ser171 substitution for Pro168 in SARS-CoV-2, which shifted the loop backbone and expanded the S4 cavity. In the SARS-CoV-2 M^pro^-nirmatrelvir complex, the P4 trifluoromethyl group is buried in a predominantly hydrophobic S4 pocket formed in part by Pro168 [46], whereas in HKU5-CoV-1 M^pro^ the expanded S4 cavity accommodates additional ordered water molecules, placing the CF_3_ group in a relatively more polar environment. Beyond nirmatrelvir, structure–activity relationship analyses and crystal structures of SARS-CoV-2 M^pro^ inhibitors have frequently shown hydrophobic P4 moieties engaging Pro168 [47-49]. This structural difference may provide opportunities for designing P4 substituents that better complement the expanded and more polar S4 environment of HKU5 Mpro.

The ensitrelvir-bound HKU5-CoV-1 M^pro^ structure closely resembles the SARS-CoV-2 M^pro^ complex [43]. Nonetheless, specific local features around the S2 and S1′ subsites deserve further scrutiny. In SARS-CoV-2 M^pro^, substitutions of Met49 in the S2-loop to Ile or Leu have been identified as resistance-associated mutations to ensitrelvir, conferring approximately one-order-of-magnitude increases in IC_50_ [43, 50]. In HKU5-CoV M^pro^, the corresponding residue is a Leu, matching the side-chain type of one of these resistance substitutions. Another key difference was found in the S1′ subsite. In SARS-CoV-2 M^pro^, Thr25 makes only limited contact with the P1′ group of ensitrelvir, whereas Met25 in HKU5-CoV M^pro^ establishes relatively stronger contacts with the P1′ indazole ring. In alignment with the functional significance of this site, it has been reported that substituting Thr25 with Ile in SARS-CoV-2 M^pro^ results in an approximately 3-fold increase in the IC_50_ for ensitrelvir [51]. Collectively, these observations indicate that, although the IC_50_ values for ensitrelvir are similar, the local environment constituted by Met25, Leu49, and Gln192 in HKU5-CoV M^pro^ (Thr25, Met49, and Gln189 in SARS-CoV-2 M^pro^) may subtly affect ensitrelvir recognition.

Beyond these local side-chain differences, the high-resolution structures provide water-mediated hydrogen bond networks of the HKU5-CoV M^pro^ active site that can be directly exploited for inhibitor optimization. In the apo enzyme, ordered water molecules occupy well-defined positions in the S1′, S1, S2, and S3–S4 subsites, forming hydrogen-bond networks with backbone and side-chain atoms. In the nirmatrelvir and ensitrelvir complexes, both inhibitors form direct hydrogen bonds that displace pre-organized waters, and additional water-mediated bridges around these substituents are observed that highlight positions where further polar extensions could be explored. Collectively, these structures provide a basis for leveraging the ordered water network in the HKU5-CoV M^pro^ active site to design optimized inhibitors that either engage or displace specific waters to enhance affinity and selectivity.

Modeling and MD simulations of HKU5-CoV-2 M^pro^ provided structural insights that, consistent with the 93% sequence identity between the two enzymes, largely preserved the inhibitor binding modes observed in the HKU5-CoV-1 crystal structures. Concurrently, this modeling work underscored the current limitations of structure prediction in antiviral design. While AlphaFold3 accurately reproduced the covalent binding pose of nirmatrelvir, it did not capture the ensitrelvir-bound conformation, particularly the inhibitor-induced inward rotation of His41 at the S2 subsite. This discrepancy suggests that, although AlphaFold3 can reliably predict global folds and the overall architecture of active sites, high-resolution experimental structures remain indispensable for structure-based drug design, particularly when precise side-chain conformations and ligand-induced rearrangements are crucial for defining inhibitor recognition [52].

Although this study advances our understanding of HKU5-CoV M^pro^ inhibition, it also has several limitations. Cell-based antiviral activity against HKU5-CoV was not evaluated in this study, so the IC_50_ values reported here should be interpreted primarily as measures of intrinsic target susceptibility rather than direct surrogates for antiviral efficacy. HKU5-CoV-1/2 M^pro^ proteins consistently exhibited approximately two-fold higher k_cat_ values than SARS-CoV-2 M^pro^, whereas K_m_ values were similar. However, the available crystal structures did not reveal an obvious structural basis for this kinetic difference, leaving the contribution of subtle active-site dynamics or product-release steps unresolved. Dissecting this mechanistic basis will require targeted mutagenesis combined with extended molecular dynamics simulations and detailed pre–steady-state and steady-state kinetic analyses.

In summary, these data define how clinically approved M^pro^ inhibitors engage a pre-emergent *Merbecovirus* HKU5-CoV protease at atomic resolution and reveal both conserved and species-specific features that can be leveraged for future drug design. By integrating high-resolution crystallography, biochemical profiling, and modeling, this work provides a framework for developing next-generation M^pro^ inhibitors with improved breadth against HKU5-related merbecoviruses and for evaluating the cross-reactive potential of existing COVID-19 M^pro^ inhibitors against emerging merbecoviruses.

## Materials and methods

### Protein expression and purification

Genes encoding the M^pro^ proteins of HKU5-CoV-1 (GenBank accession: YP_001039961.1), HKU5-CoV-2 (Genbase accession number: C_AA085189.1), and SARS-CoV-2 (GenBank accession: WPE12197.1) were codon-optimized for *Escherichia coli* expression and synthesized. These genes were subcloned into the pET28a(+) vector, which contained an N-terminal His_6_–small ubiquitin-related modifier (SUMO) affinity tag, and recombinant plasmids were transformed into *E. coli* BL21(DE3) CodonPlus RIL cells. Cultures were grown in Luria–Bertani media containing 50 μg/mL kanamycin at 37°C. When OD_600_ reached 0.5-0.6, protein expression was induced with 0.5 mM isopropyl β-D-1-thiogalactopyranoside, followed by incubation at 18°C for 16 h. The cells were harvested and stored at -80°C.

For purification, cell pellets were resuspended in a lysis buffer containing 20 mM Tris-HCl (pH 8.0), 150 mM NaCl, and 2 mM β-mercaptoethanol and disrupted *via* sonication on ice. After centrifugation (19,000 × *g*, 4°C, 30 min), the supernatant was loaded onto a Ni-NTA affinity column (Cytiva). The impurities were washed with lysis buffer supplemented with 20 mM imidazole and His_6_-SUMO-tagged proteases were eluted with the lysis buffer supplemented with 200 mM imidazole. The His_6_-SUMO tag was removed by Ulp1 protease and applied to HiTrap Capto Q ImpRes column (Cytiva). Proteins were eluted with a linear NaCl gradient (0-1 M) and further purified by size-exclusion chromatography on a Superdex 200 16/600 column (Cytiva) equilibrated with SEC buffer containing 20 mM HEPES (pH 7.5), 150 mM NaCl, and 1 mM tris(2-carboxyethyl)phosphine. The purified proteins were concentrated using Vivaspin 20 (Sartorius) centrifugal filters and stored at -80°C.

### Enzyme kinetics assays

For kinetic assays, reactions were carried out in assay buffer consisting of 25 mM HEPES (pH 7.5), 150 mM NaCl, 1 mM EDTA, 1 mM dithiothreitol, and 5% (v/v) dimethyl sulfoxide. HKU5-CoV-1, HKU5-CoV-2, and SARS-CoV-2 M^pro^ proteins were used at a final enzyme concentration of 100 nM. A fluorogenic FRET peptide

(Dabcyl-KTSAVLQ↓SGFRKME-Edans; MedChemExpress) was employed as the substrate at final concentrations ranging from 1.25 to 160 μM. Reactions (100 μL) were initiated by mixing enzyme and substrate in black 96-well plates, and fluorescence was monitored at 340/460 nm (excitation/emission) with a 425 nm cutoff at 25°C. Initial velocities were fitted to the Michaelis–Menten equation to determine K_m_ and k_cat_. The kinetic parameters are reported as mean ± SD from three independent experiments (each performed in triplicate). All analyses were carried out using GraphPad Prism 10 software (Boston, MA, USA).

### Enzymatic inhibitory assays

Enzymatic inhibitory assays were performed at 25°C using recombinant HKU5-CoV-1, HKU5-CoV-2, and SARS-CoV-2 M^pro^ proteins (100 nM) in assay buffers. Nirmatrelvir (PF-07321332; MCE, cat. no. HY-138687) and ensitrelvir (S-217622; MCE, cat. no. HY-143216) were prepared as 50 mM stock solutions in DMSO and serially diluted to final concentrations ranging from 0.0001 to 100 μM. The enzymes and inhibitors were pre-incubated at 25°C for 15 min in the assay buffer. Reactions were initiated by adding the substrate (final concentration of 20 μM) to opaque 96-well black-bottom plates (total volume of 100 μL per well). Fluorescence was measured at an excitation/emission wavelength of 340/460 nm with a 425 nm cutoff. IC_50_ values were determined by fitting dose–response curves using GraphPad Prism 10 (GraphPad Software, Boston, MA, USA). Experiments were performed as three independent biological replicates with triplicate wells per concentration, and IC_50_ values are reported as mean ± SD.

### Crystallization and X-ray diffraction data collection

The purified HKU5-CoV-1 M^pro^ was concentrated to 16 mg/mL in SEC buffer for crystallization. For inhibitor-bound complexes, the protein was pre-incubated with 1 mM nirmatrelvir or ensitrelvir before crystallization trials. Apo and complex crystals were obtained under similar conditions and further optimized using the hanging-drop vapor diffusion method at 14°C. The optimized reservoir solution contained 0.16 M MgCl_2_, 0.08 M Tris-HCl (pH 8.5), 28% (w/v) PEG 4000, and 20% (v/v) glycerol. The crystals were flash-cooled in liquid nitrogen stream without additional cryoprotection. X-ray diffraction data were collected at beamline 11C of the Pohang Accelerator Laboratory at a wavelength of 0.97942 Å [53].

### Data processing and structure determination

The diffraction data were indexed, integrated, and scaled using the HKL-2000 program [54]. The apo HKU5-CoV-1 M^pro^ structure was solved by molecular replacement (MR) using Phaser in the Phenix suite [55]. An AlphaFold3-generated model of the HKU5-CoV-1 M^pro^ served as the initial search template [44, 56], and the structures of the inhibitor-bound complexes were solved by MR using the refined apo structure as the search model. Model building was performed through iterative cycles of manual adjustment in Coot, followed by refinement with phenix.refine [57, 58].

### Computational modeling of the HKU5-CoV-2 M^pro^–inhibitor complexes using MD simulations

MD simulations of the HKU5-CoV-2 M^pro^–inhibitor complex models were performed using the GROMACS software [59]. System preparation and input file generation were conducted using the CHARMM-GUI solution builder with the CHARMM36m force field [60-62]. Each protein was solvated in a cubic box of TIP3P water molecules, and the system was neutralized and adjusted to a salt concentration of 150 mM KCl. Energy minimization was performed with position restraints on protein atoms using the steepest-descent integrator, followed by 100 ps NVT and 500 ps NPT equilibration while maintaining the position restraints. MD simulations were performed under NPT conditions at 303.15 K and 1.0 bar using a time step of 0.002 ps. Temperature and pressure were controlled using the v-rescale thermostat and C-rescale barostat, respectively. Van der Waals interactions were calculated using a cutoff scheme (rvdw = 1.2 nm), while electrostatic interactions were treated using the particle mesh Ewald method. Hydrogen bonds were constrained using the LINCS algorithm, and trajectory analyses were performed using the GROMACS tools after correcting for periodic boundary conditions and centering the protein to remove translational drift.

### Accession number

The atomic coordinates and structure factors have been deposited in the Protein Data Bank (PDB) under accession codes 9VVC (apo), 9VVF (nirmatrelvir-bound), and 9WM7 (ensitrelvir-bound).

## CRediT authorship contribution statement

**Hyojin Kim:** Writing – review & editing, Writing – original draft, Investigation, Visualization, Formal analysis, Data curation, Conceptualization. **Jinsook Ahn:** Writing – review & editing, Visualization, Validation, Software. **Juyeon Lee:** Methodology, Supervision. **Seungheon Jung:** Investigation, Resources. **Ji Woo Kim:** Investigation, Resources. **Byungil Kim:** Conceptualization, **Nam-Chul Ha:** Supervision, Project administration, Writing – review & editing, Conceptualization. **Inseong Jo:** Writing – review & editing, Writing – original draft, Supervision, Project administration, Methodology, Funding acquisition, Conceptualization.

## Acknowledgments

This work was supported by the National Research Foundation of Korea (NRF) grant funded by the government of Korea (MSIT) (RS-2024-00401962 and RS-2026-25474560 to IJ), and by intramural funds from the Korea Research Institute of Chemical Technology (KK2633-10 to IJ). We thank the staff at the 11C beamline of Pohang Accelerator Laboratory (PAL) for their support during the X-ray data collection.

## Data availability

Data will be made available on request.

